# Unbiased Age-Appropriate Structural Brain Atlases for Chinese Pediatrics

**DOI:** 10.1101/385211

**Authors:** Tengda Zhao, Xuhong Liao, Vladimir S. Fonov, Weiwei Men, Yanpei Wang, Shaozheng Qin, Shuping Tan, Jia-Hong Gao, Alan Evans, Sha Tao, Qi Dong, Yong He

**Author notes:** Corresponding author: Yong He, Ph.D., State Key Laboratory of Cognitive Neuroscience and Learning, Beijing Key Laboratory of Brain Imaging and Connectomics, IDG/McGovern Institute for Brain Research, Beijing Normal University, Beijing, 100875, China.

## Abstract

In magnetic resonance imaging (MRI) studies of children brain development, structural brain atlases usually serve as important references of pediatric population in which individual images are spatially normalized into a common or standard stereotactic space. However, the existing popular children brain atlases (e.g., National Institutes of Health pediatric atlases, NIH-PD atlases) are made mostly based on MR images from Western populations, and are thus insufficient to characterize the brains of Chinese children due to the neuroanatomical differences that are relevant to genetic and environmental factors. By collecting high-quality T1- and T2- weighted MR images from 328 typically developing Chinese children aged from 6 to 12 years old, we created a set of age-appropriate Chinese pediatric (CHN-PD) atlases using an unbiased template construction algorithm. The CHN-PD atlases included the *head/brain* templates, the symmetric brain template, the gender-specific brain templates and the corresponding tissue probability atlases. Moreover, the atlases contained multiple age-specific templates with a one-year interval. A direct comparison of the CHN-PD and the NIH-PD atlases revealed remarkable anatomical differences bilaterally in the lateral frontal and parietal regions and somatosensory cortex. While applying the CHN-PD atlases to two independent Chinese pediatric datasets (N = 114 and N = 71, respectively), machine-learning regression approaches revealed higher prediction accuracy on brain ages than the usage of NIH-PD atlases. These results suggest that the CHN-PD brain atlases are necessary and important for future typical and atypical developmental studies in Chinese pediatric population. Currently, the CHN-PD atlases have been released on the NITRC website (https://www.nitrc.org/projects/chn-pd).

## Introduction

Modern advances in multi-modal magnetic resonance imaging (MRI) offer an unprecedented opportunity to explore the structural and functional development of the pediatric brain *in vivo*. A typical research framework is achieved by normalizing individual brain images into a common or standard stereotactic space using a prior structural brain atlas as a reference (Ashburner and Friston, 1999; Collins, et al., 1998; Smith, et al., 2004), such as the International Consortium for Brain Mapping templates (ICBM152 templates) at the Montreal Neurological Institute (MNI) space (Evans, et al., 2012; Lancaster, et al., 2007). Due to the rapid development of the brain, structural brain atlases specific to young children have been generated for pediatric MRI investigations (Fonov, et al., 2011; Luo, et al., 2014; Oishi, et al., 2018; Richards, et al., 2016; Sanchez, et al., 2012; Wilke, et al., 2008; Wilke, et al., 2002; Xie, et al., 2015). It has been argued that adopting such age-appropriate brain templates in pediatric participants can reduce the requirement for spatial deformation during image normalization and maintain more pediatric characteristics of individual brain such as a thicker cerebral cortex compared with adult templates (Fonov, et al., 2011; Yoon, et al., 2009). However, the existing children brain templates are constructed mostly based on Western pediatric populations (Fonov, et al., 2011; Oishi, et al., 2018; Richards, et al., 2016; Sanchez, et al., 2012; Wilke, et al., 2008; Wilke, et al., 2002), with a typical case being the widely used National Institutes of Health pediatric atlases (NIH-PD) (Fonov, et al., 2011). These existing brain templates are not ideal for use in Chinese pediatrics studies (Richards and Xie, 2015), since Chinese adults and children have unique neuroanatomical features in the brain size and shape as compared to Western people (Bai, et al., 2012; Liang, et al., 2015; Tang, et al., 2010; Tang, et al., 2018; Xie, et al., 2015). Different growth trajectories of some brain structures between Chinese and North American children have also been reported (Guo, et al., 2007; Xie, et al., 2014). To make an accurate brain representation of Chinese pediatric population, it would be necessary and important to create age-specific atlases based on the MR images of Chinese children.

When constructing pediatric brain atlases, two common factors need be considered: brain asymmetry and the gender effect. The development of child brain is asymmetric or lateralized in both structure and function (Agcaoglu, et al., 2015; Song, et al., 2014; Zhong, et al., 2016; Zhou, et al., 2013), which is related to the specialization of language and motor functions and may underlie developmental brain disease phenotypes (Herbert, et al., 2002; Shaw, et al., 2009; Toga and Thompson, 2003). To estimate the development of brain asymmetry, symmetric brain models are expected to treat both hemispheres equally. Symmetric brain templates have been created for Western pediatric populations (Fonov, et al., 2011). However, Chinese children may have different brain asymmetries from Western children due to the genetic and cultural factors. Studies have revealed that the visual process of Chinese characters engages more bilateral temporo-occipital regions of the brain than alphabetic languages (Cao, et al., 2009; Mei, et al., 2015; Xue, et al., 2005). Developmental brain disorders with asymmetric abnormalities such as dyslexia (Beaton, 1997; Leonard and Eckert, 2008) also present unique brain disruptions in Chinese children (Siok, et al., 2008; Siok, et al., 2004). The construction of symmetric brain templates for Chinese children is important for further investigations of the Chinese pediatrics. Equally important, previous MRI studies have revealed gender-specific differences in the brain anatomy of typical and atypical developmental populations (De Bellis, et al., 2001; Evans, et al., 2014; Gennatas, et al., 2017; Good, et al., 2001; Peper, et al., 2011). Several developmental disorders, such as autism spectrum disorder or attention deficit hyperactivity disorder (ADHD), show a gender-specific prevalence and symptomatology (Vértes and Bullmore, 2015). In an extreme case, sex chromosome related developmental disorders occur only in a single-sex population (Cutter, et al., 2006; Murphy, et al., 1993). Under these situations, a gender-specific brain template can attain a more accurate characterization of the pediatric brain than that of a general population.

To date, there are only two previous MRI studies towards the construction of Chinese pediatric brain templates (Luo, et al., 2014; Xie, et al., 2015). Specifically, Luo, et al. (2014) built a single brain template within a narrow age range of 5 to 8 years old based on structural MR images of 53 Chinese children. Xie, et al. (2015) generated a series of pediatric brain templates based on structural MR images of 138 Chinese children within an age range of 8 to 16 years old with a 2-year interval. However, the application of the two brain templates is still limited for Chinese pediatric studies due to several methodological issues as follows. First, the quality of these two brain templates is inadequate due to the low signal to noise ratio of the 1.5T MRI scanner (Luo, et al., 2014) and the relatively small sample size (Luo, et al., 2014; Xie, et al., 2015). Moreover, neither study provided symmetric and gender-specific templates. Second, both studies employed customized coordinates that are different from the popular ICBM152 and NIH-PD templates at the MNI space (Fonov, et al., 2011). Third, it needs to note that these two previous studies revealed only overall differences in the brain circumference and the deformation cost during registration between Chinese and Western brain templates (Liang, et al., 2015; Tang, et al., 2010; Xie, et al., 2015). Considering spatially distributed variations of the brain regions that could be derived from genetic and environmental effects during development, it is important to examine the detailed regional anatomical differences between Chinese and Western pediatric brain templates.

In the present study, we aimed to create a set of high quality Chinese pediatric (CHN-PD) atlases for school-aged children (6-12 years old). To do this, we first collected high-quality MRI images of a large sample (328 participants) at the state-of-the-art 3T Siemens Prisma scanner. Then, we employed an unbiased template construction algorithm to generate the average *head/brain,* symmetric and gender-specific MRI templates with finer one-year age intervals. Finally, to further investigate the necessity and applicability of the proposed CHN-PD atlases, we assessed the regional difference between the CHN-PD and the NIH-PD templates and evaluated their prediction power on brain age in two independent Chinese pediatric datasets (N = 114 and N = 71, separately) when using different brain templates during the spatial normalization.

## Materials and Methods

### Participants

This study included three datasets of healthy Chinese children (Table 1): i) a principal dataset (Dataset 1) of 328 participants aged from 6–12 years old (9.03 ± 1.36) scanned at the Peking University (PKU), ii) an independent dataset (Dataset 2) of 114 participants aged from 6–12 years old (9.06 ± 1.38) scanned at the Beijing HuiLongGuan (HLG) hospital in China, and iii) another independent public dataset (Dataset 3) including 71 participants aged from 8 to 12 years old (10.26 ± 1.78) that were obtained from the Beijing site of the ADHD200 dataset via the International Data-sharing Initiative (Consortium, 2012; Fair, et al., 2012). Participants of Datasets 1 and 2 were recruited from local primary schools in Beijing and written informed consent was obtained from the parents/guardians of the children. Detailed information about the participants in Dataset 3 can be seen at the public data sharing website (http://fcon_1000.projects.nitrc.org/indi/adhd200/). All these MRI scans have passed through a strict quality control criterion, before which the initial data collection included 359 participants in Dataset 1, 131 participants in Dataset 2 and 71 participants in Dataset 3. The detailed quality control procedure is as follows: i) all images were first reviewed by an experienced neurologist to exclude neurological abnormalities; ii) careful visual inspections with a scan rating procedure were conducted by two experienced raters separately, similar to the protocol used in the human connectome project (HCP) (Marcus, et al., 2013); iii) images that were assigned a quality of better than fair by both raters were retained. This study was approved by the ethics committee of Beijing Normal University. Notably, Dataset 1 was used for the construction of the Chinese age-appropriate MRI templates, and Dataset 2 and 3 were used for the evaluation of the template effect on age prediction. The age and gender distributions of these samples are listed in Table 1.

**Table 1.** Demographic information for the three datasets

|  | Age range (years) | Number | Gender (girl/boy) |
| --- | --- | --- | --- |
| Dataset 1 (PKU site) | 6-12 ($9.03 \pm 1.36$) | 328 | 153/175 |
| Dataset 2 (HLG site) | 6-12 ($9.06 \pm 1.38$) | 114 | 47/67 |
| Dataset 3 (Beijing site from ADHD200) | 8-12 ($10.26 \pm 1.78$) | 71 | 46/25 |

PKU, Peking University; HLG, HuiLongGuan hospital

### Image acquisition

In Datasets 1 and 2, high quality T1- and T2-weighted images were acquired for each participant on 3T Siemens Prisma scanners. The detailed scanning parameters are as follows: T1 weighted images: repetition time (TR) = 2530ms, echo time (TE) = 2.98ms, inversion time (TI) = 1100ms, flip angle (FA)= 7°, acquisition matrix = 256 × 256, field of view (FOV) = 256 × 224 mm^2^, slices = 192, slice thickness = 1mm, BW = 240Hz/Px; T2 weighted images: TR = 3200ms, TE = 564ms, acquisition matrix = 320 × 320, FOV = 224 × 224 mm^2^, slices = 256, slice thickness = 0.7mm, BW = 744Hz/Px. In dataset 3, images were acquired using a 3T Siemens Trio scanner, and the scanning parameters are as follows: T1-weighted magnetization-prepared rapid acquisition gradient echo sequences, 128 slices, slice thickness = 1.33 mm, TR = 2530 ms, TE = 3.39 ms, TI = 1100 ms, FA = 7°, acquisition matrix = 256 × 256, FOV = 256 × 256 mm^2^, average =1.

### Data preprocessing

All MRI scans were preprocessed as follows (Fig. 1, left panel): i) the intensity inhomogeneity of each scan was corrected using N4 correction (Tustison, et al., 2010); ii) brain masks were created using the robust BET estimation from the FSL (FMRIB Software Library), and skull outlines were generated using the BET2 command (Smith, 2002); iii) a hierarchical linear registration (nine-parameter affine transformation) of each scan into the ICBM152 linear brain template was conducted using the Revised BestLinReg algorithm (Dadar, et al., 2018); iv) the image intensity was scaled to the same range of the template resulting in intensities between 0 and 100 (Nyúl and Udupa, 1999); and v) tissue segmentations were implemented using the well-validated CIVET 2.1 pipeline to obtain the probability maps of gray matter (GM), white matter (WM) and cerebral spinal fluid (CSF) of each individual (http://www.bic.mni.mcgill.ca/ServicesSoftware/CIVET-2-1-0-Table-of-Contents).

**Figure 1.**
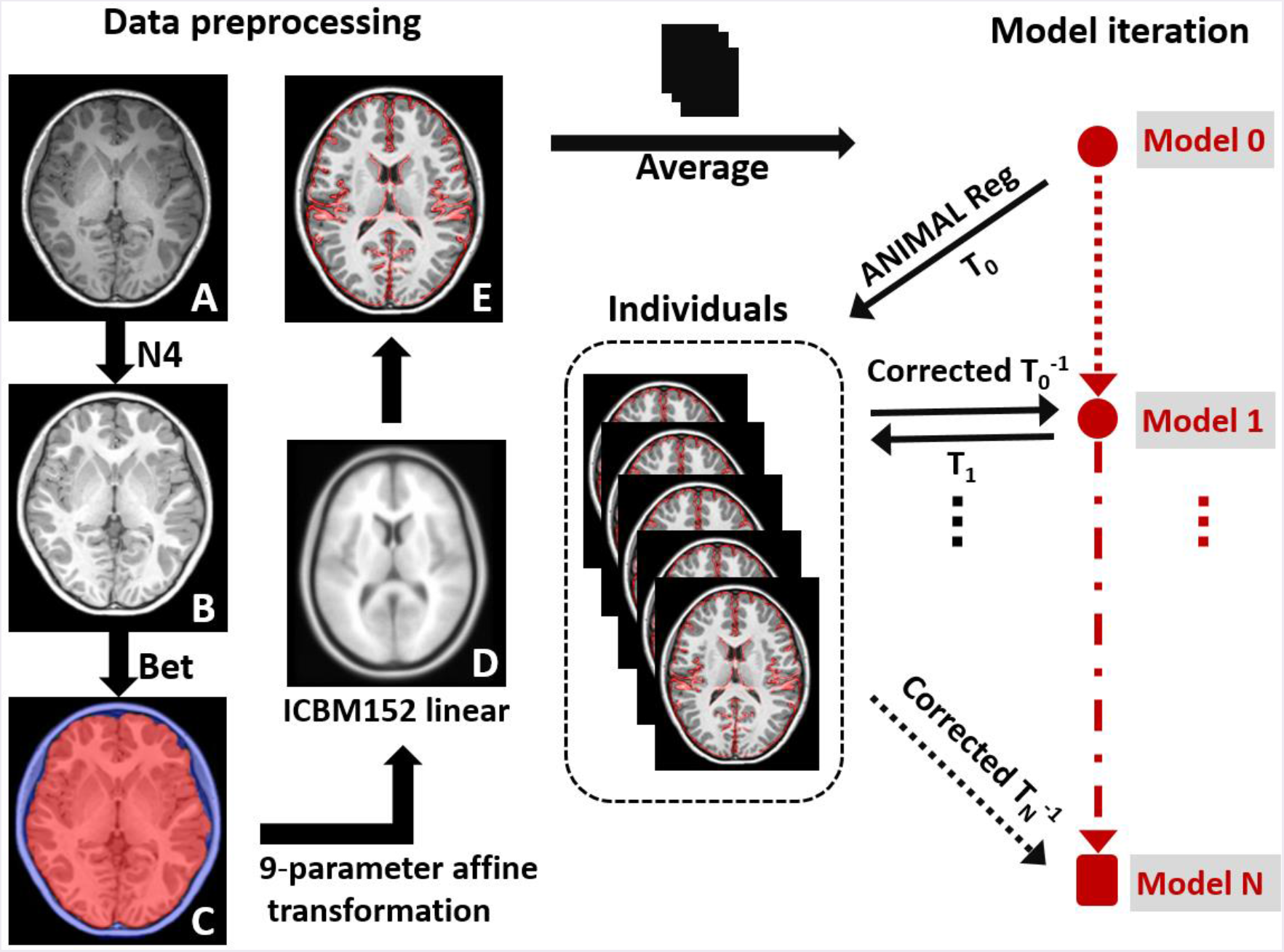
The flowchart shows the data preprocessing and template iteration process. The individual MRI scan (A) was first corrected for intensity inhomogeneity by using N4 correction (B). The brain mask and skull outlines (C) were created using the robust BET estimation from the FSL; A hierarchical linear (nine-parameter affine transformation) registration of the individual image to the ICBM152 linear template (D) was conducted using the Revised BestLinReg algorithm, and the image intensity was scaled to the same range as the target (D). The resulting individual images (E) were averaged to generate the initial template (Model 0) and used to generate individual transformations (T_0_) via ANIMAL nonlinear registration. This deformation (T_0_) was further corrected by the average transformation and applied to individual images (corrected T_0_^−1^) to create the new averaging approximation. This iterative process continued until convergence was reached.

### Template construction

The unbiased template construction algorithm adopted in our study was proposed by Fonov and colleagues (Fonov, et al., 2011) based on previous works (Guimond, et al., 1998; Guimond, et al., 2001). This procedure has been widely applied to generate MRI templates including the ICBM152 brain template, the NIH-PD brain template and the standard spinal cord template (De Leener, et al., 2018; Fonov, et al., 2011; Fonov, et al., 2009; Fonov, et al., 2014). This iterative construction algorithm can capture both the average intensity and the average shape of the brain at a population level. We listed its brief process as follows (Fig. 1, right panel). First, the ICBM152 linear brain template was used as the initial reference target template onto which each preprocessed T1-weighted image was mapped nonlinearly. The generated individual transformations were further corrected to remove the bias via the averaging transformation. The new approximation was then generated by applying each corrected transformation to the corresponding individual T1-weighted image and then averaging all images together. The above iteration continued until convergence was reached. During each iteration step, a nonlinear registration of Automatic Nonlinear Image Matching and Anatomical Labeling (ANIMAL) (Collins, et al., 1995) was performed with an increasingly finer grid step size and blurring kernel. We adopted the following hierarchical schedule as the NIH pediatric brain template (Fonov, et al., 2011): 4 iterations at 32 mm resolution, 4 iterations at 16 mm, 4 iterations at 8 mm resolution, 4 iterations at 4 mm, and 4 iterations at 2 mm, which yielded a progressively accurate average template. The T2-weighted brain templates and tissue probability atlases were created by warping their corresponding preprocessed images of each individual based on the final deformation field for the creation of the T1-weighted brain template and then generating an average image, separately.

Using the above template construction framework, we generated a set of Chinese pediatric templates based on the PKU dataset (Dataset 1): i) the whole-head and brain-extracted MRI volumes were performed separately during the model iteration to generate both *head/brain* templates; ii) the symmetric brain templates were generated by including left-right flipped scans of each participant and enforcing symmetric deformation fields (Fonov, et al., 2011); iii) the gender-specific brain templates were generated by using the MRI volumes of male or female participants separately; iv) to generate the MRI templates specific to each age sub-range, we divided all children participants into six age subgroups with one-year intervals (see Table 2 for detailed information about the participants in each age subgroup) and same construction procedures were conducted for each group.

**Table 2.**
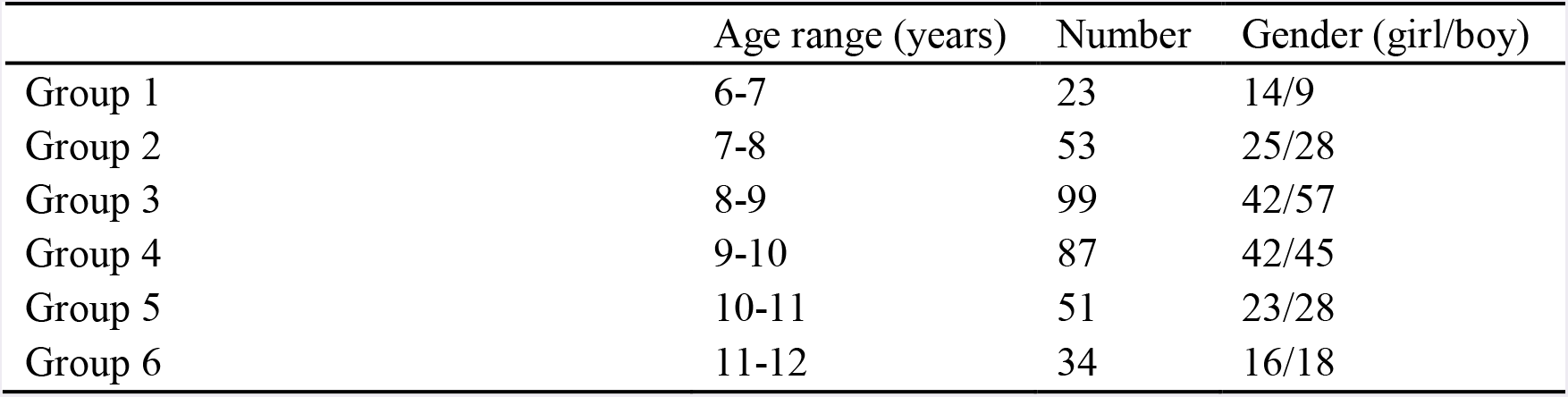
Demographic information for each age subgroup of Dataset 1

|  | Age range (years) | Number | Gender (girl/boy) |
| --- | --- | --- | --- |
| Group 1 | 6-7 | 23 | 14/9 |
| Group 2 | 7-8 | 53 | 25/28 |
| Group 3 | 8-9 | 99 | 42/57 |
| Group 4 | 9-10 | 87 | 42/45 |
| Group 5 | 10-11 | 51 | 23/28 |
| Group 6 | 11-12 | 34 | 16/18 |

### Evaluation of the anatomical difference in brain regions between the CHN-PD and NIH-PD templates

When estimating the regional difference between the proposed CHN-PD template and the commonly used NIH-PD template, we extracted a subgroup of PKU children (Dataset 1) with the same age and gender distribution and the same number of subjects as those used for generating the NIH-PD template at 7–11 years old to reduce the effect of sample distribution. Using this subgroup data, a new Chinese pediatric brain template was created and the NIH-PD brain template was co-registered into this template with a linear nine-parameter affine transformation.

Instead of the usual visual inspections, we estimated the anatomical differences between these two brain templates by calculating two quantitative image indexes for each brain region. Specifically, the Brodmann atlas was first adopted to parcellate each brain template into 82 regions. For each region, we calculated the mean square difference (MSD) (Holden, et al., 2000) of the gray matter probability maps to assess the absolute differences in the gray matter morphology and the normalized cross correlation (NCC) (Zhao, et al., 2006) across voxels of the T1-weighted brain templates to reflect the spatial similarity of the anatomical structures. The detailed definition is as follows:

For a given region μ in template A and template B, the mean square difference was defined as:

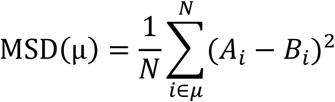

where *i* is the *i*th voxel of region μ, and *A_i_* or *B_i_* is the value of the voxel *i* of template A or B, respectively, which refers to the probability of GM in the current definition. The normalized cross correlation was defined as follows:

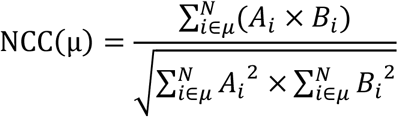

where *i* is the ith voxel of region μ, and *A_i_* or *B_i_* is the value of the voxel *i* of template A or B, respectively, which refers to the intensity of the T1-weighted brain template in the current definition. The MSD shows the mean magnitude of the voxel-wise absolute difference between two template regions, while the NCC is commonly used as the cost function during image matching and indicates the spatial similarity of two brain regions. Here, we used the index of 1-NCC to present the anatomical difference of each region between brain templates. The calculation of the NCC was conducted with the Statistical Parametric Mapping (SPM) software (Friston, et al., 1994) (https://www.fil.ion.ucl.ac.uk/spm/).

### Evaluation of the template effect on the accuracy of age prediction

To determine whether the proposed CHN-PD atlases have advantages over Western pediatric brain atlases in Chinese children studies, we performed a comparative analysis between the CHN-PD and NIN-PD brain templates (7–11 years old, sample distribution matched) on the prediction power of brain age. Specifically, to reduce the over-fitting effect, we adopted two independent datasets to train and test the prediction model of brain age, separately. First, the HLG child subjects (Dataset 2) were used as the training sample and another independent set of healthy samples in the ADHD200 dataset (Dataset 3) was used as the testing sample. For each subject, the T1-weighted brain image was separately normalized to the new Chinese children brain template and to the NIH-PD brain template at 7–11 years old using the hierarchical ANIMAL nonlinear registration. The Brodmann mask was warped separately into two brain templates by a non-linear transformation to locate feature voxels for the prediction model. We implemented two most widely used machine-learning regression strategies (Cui and Gong, 2018; Drucker, et al., 1997; Tipping, 2001) to predict the individual age based on the normalized T1-weighted brain images. The first strategy is a support-vector regression (SVR) model with a linear kernel function (Drucker, et al., 1997). Common settings with C = 1 and epsilon = 0.001 were adopted. The second strategy is the relevance vector regression (RVR), which is a Bayesian-formulated regression framework (Tipping, 2001). After model training and testing, the Pearson correlation coefficient between the actual and predicted ages was calculated to assess the prediction accuracy. To reduce the effect of the feature preprocessing methods, we applied three feature preprocessing approaches, including the raw features (untreated), scaling and normalization, and then repeated the whole prediction procedure. Finally, we exchanged the training and test samples to make a cross validation of the prediction. For the code implementation, the LIBSVM function was used in the SVR model (https://www.csie.ntu.edu.tw/~cjlin/libsvm/) (Chang and Lin, 2011), and the PRoNTo toolbox (http://www.mlnl.cs.ucl.ac.uk/pronto/) was used in the RVR model (Schrouff, et al., 2013).

## Results

### Convergence of the template construction algorithm in Chinese pediatric atlases

In this study, all types of MRI templates were constructed through the hierarchical model iteration processes, and qualitative progression was observed along with the iterations. Figure 2A and B illustrate a detailed view of the intermediate models during the construction of the brain template over the full age range (6–12 years old). We showed the voxel-wise standard deviation map across the individual scans and the averaged models at different iterations and resolution steps. The standard deviation at the voxel level was reduced in every four iterations, and the anatomical details became distinct gradually around the neighboring voxels, showing a successful convergence (Fig. 2A and B). The standard T1-weighted *brain* template was generated at the 20^th^ iteration. The root mean square (RMS) of the intensity standard deviation and biases of the average deformation in each iteration step were plotted for all types of MRI templates (Fig. 2C). As the procedure advanced, the RMS decreased progressively across iterations for the averaging models, indicating that the optimization procedure was reaching a minimum. Similar convergence processes were found for the averaged *head/brain,* symmetric, and gender-specific templates (Fig. 2C, left panel) and also for each age subgroup templates (Fig. 2C, right panel).

**Figure 2.**
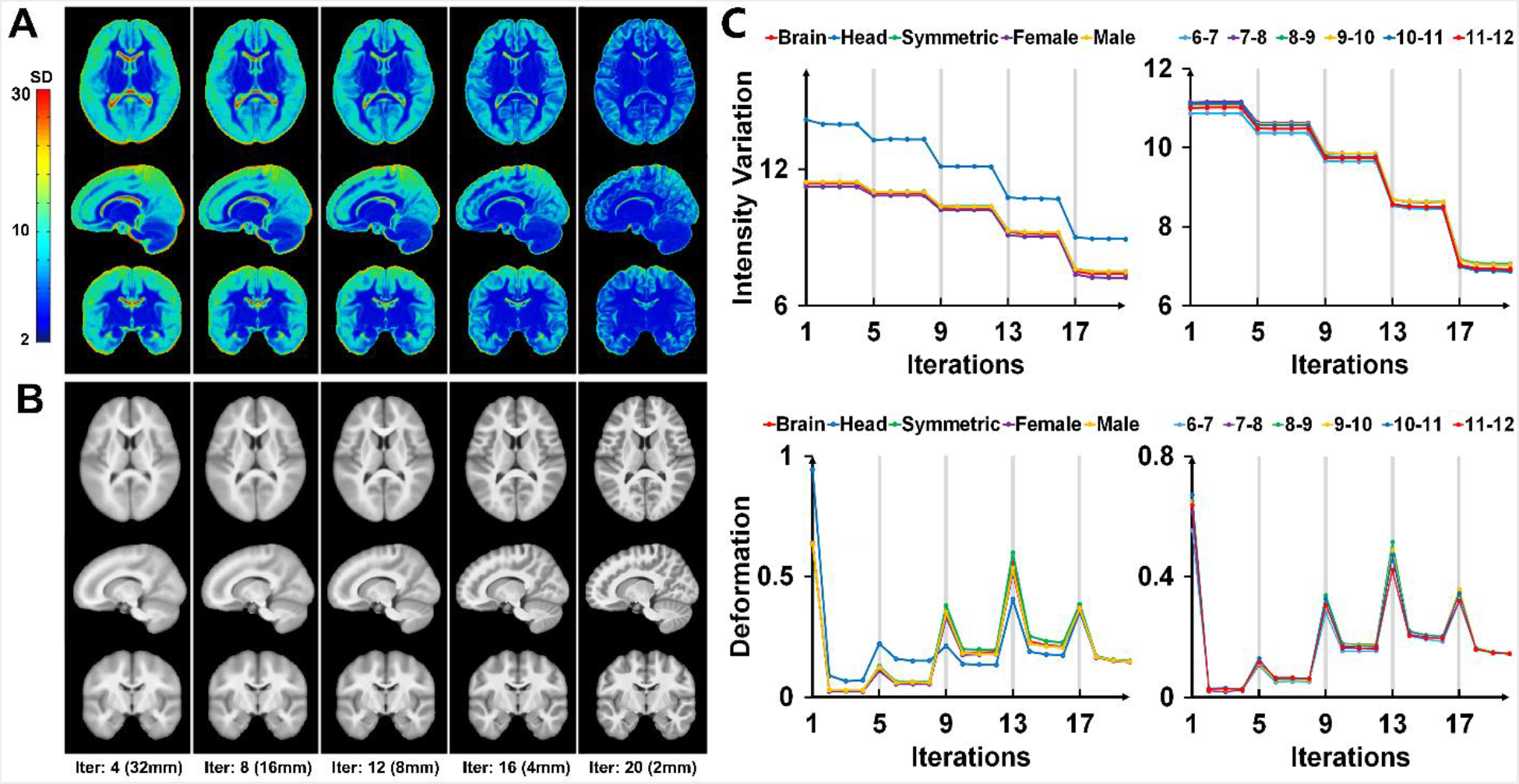
The average MRI templates generated in the hierarchical matching process. (A) Color scale images show the intensity standard deviation (SD) of each averaged brain model at the certain level of step size during the iteration. (B) Gray scale images show the average T1-weighted brain models at the corresponding steps. (C) The root mean square of the intensity SD (upper panel) and the biases in the average deformation (below panel) at each iteration are given for the full age range (left panel) and age sub-ranges (right panel) MRI templates. The gray line shows the different blurring kernels during the iterations.

### Averaged structural MRI atlases for Chinese pediatrics (CHN-PD atlases)

Figure 3 shows the Chinese pediatric MRI templates over the full age range (6–12 years old), where they are shown in axial, sagittal and coronal views, separately. The first five columns show the detailed slices of the *head/brain,* symmetric and gender-specific T1-weighted template, and the subsequent columns show the slices of the T2-weighted brain template and the tissue probability atlases of the GM, WM, and CSF. In each slice, these templates exhibit distinct anatomical structures in the cerebral and subcortical regions, cerebellum, and brainstem. Figure 4 shows the detailed age-specific templates at one-year intervals, including the T1-weighted brain templates and the standard tissue probability atlases. By visual inspection, particularly similar spatial locations of the brain gyri and sulci were found in each template and slight anatomical changes were found in the peripheral gray and white matter junctions. All of these templates are publicly available in NIFTI format in the Chinese pediatric atlases (CHN-PD atlas) project on the NITRC website (https://www.nitrc.org/projects/chn-pd/).

**Figure 3.**
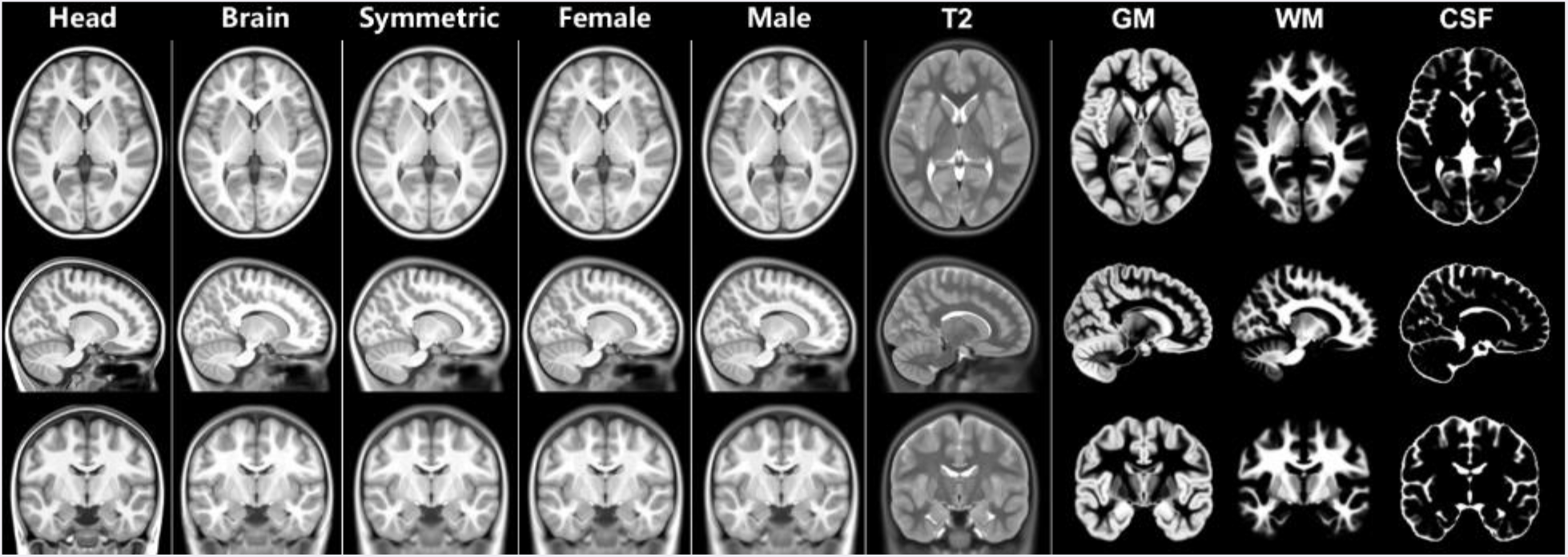
The detailed slices of the average T1-weighted *head/brain* templates, the symmetric template and the gender-specific templates *female/male*) and the slice view of the average T2-weighted brain templates and the tissue probability templates in Chinese pediatric atlases (6-12 years old).

**Figure 4.**
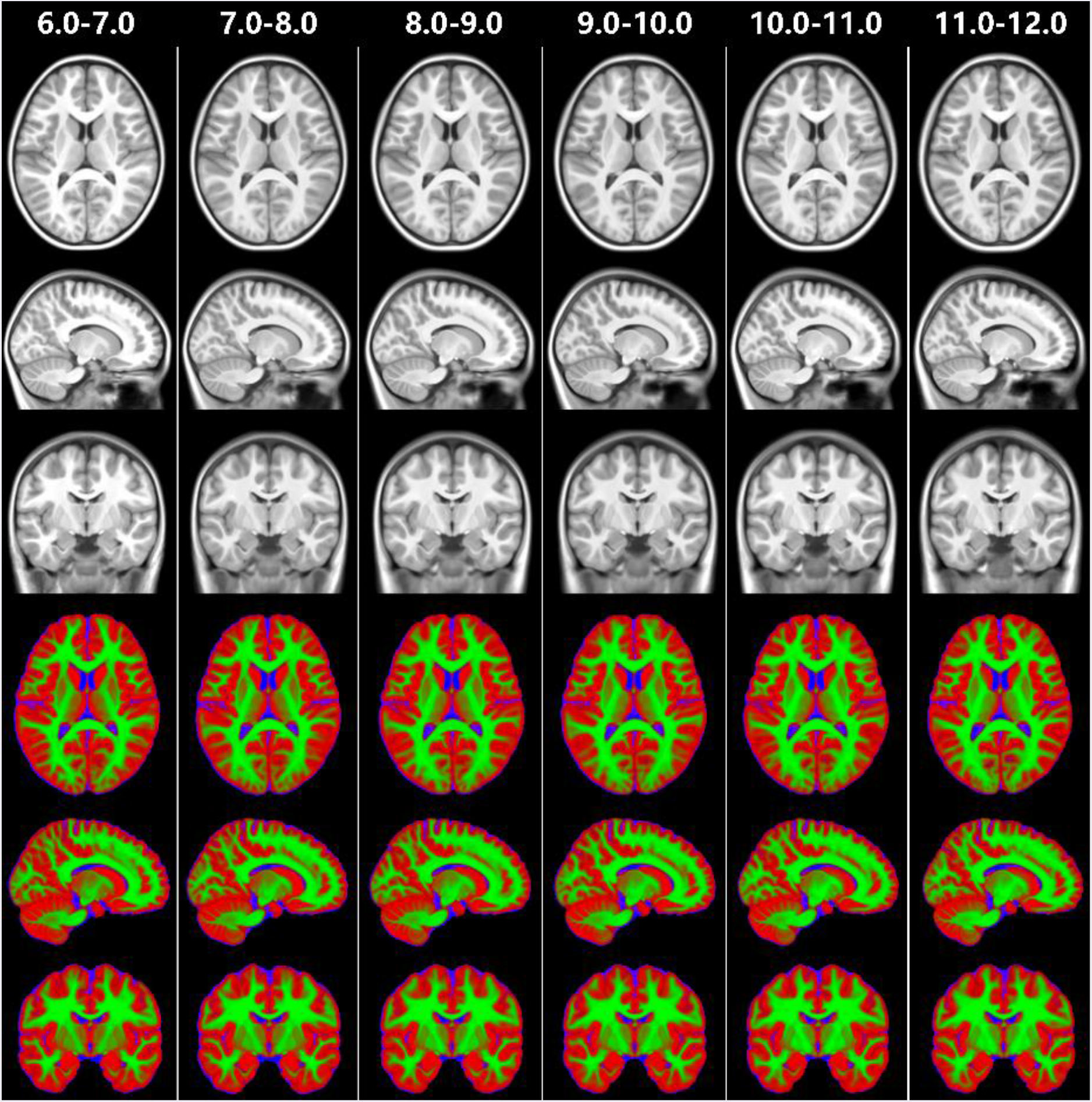
The detailed slices of the average T1-weighted brain templates (upper panel) and the combined tissue class atlas (below panel) of the age sub-groups with one-year intervals. Colors in red represents gray matter, green represents white matter, and blue represents CSF.

### Regional differences between the CHN-PD and the NIH-PD atlases

From the perspective of the absolute differences in the gray matter probability maps between two brain templates, regions showing relatively high anatomical differences were mainly located in bilateral angular gyrus and supramarginal gyrus (parts of Wernicke’s area), bilateral dorsolateral prefrontal cortex and inferior frontal regions (including Broca’s area), and bilateral somatosensory cortex (Fig. 5, upper panel, and Table 3). From the perspective of the voxel-wise spatial similarity, a consistent distribution of regional differences was found, which showed high structural differences in the bilateral angular gyrus and dorsolateral and inferior frontal gyrus between the Chinese pediatric atlases (CHN-PD) and NIH pediatric brain template (NIH-PD) (Fig. 5, lower panel, and Table 3).

**Figure 5.**
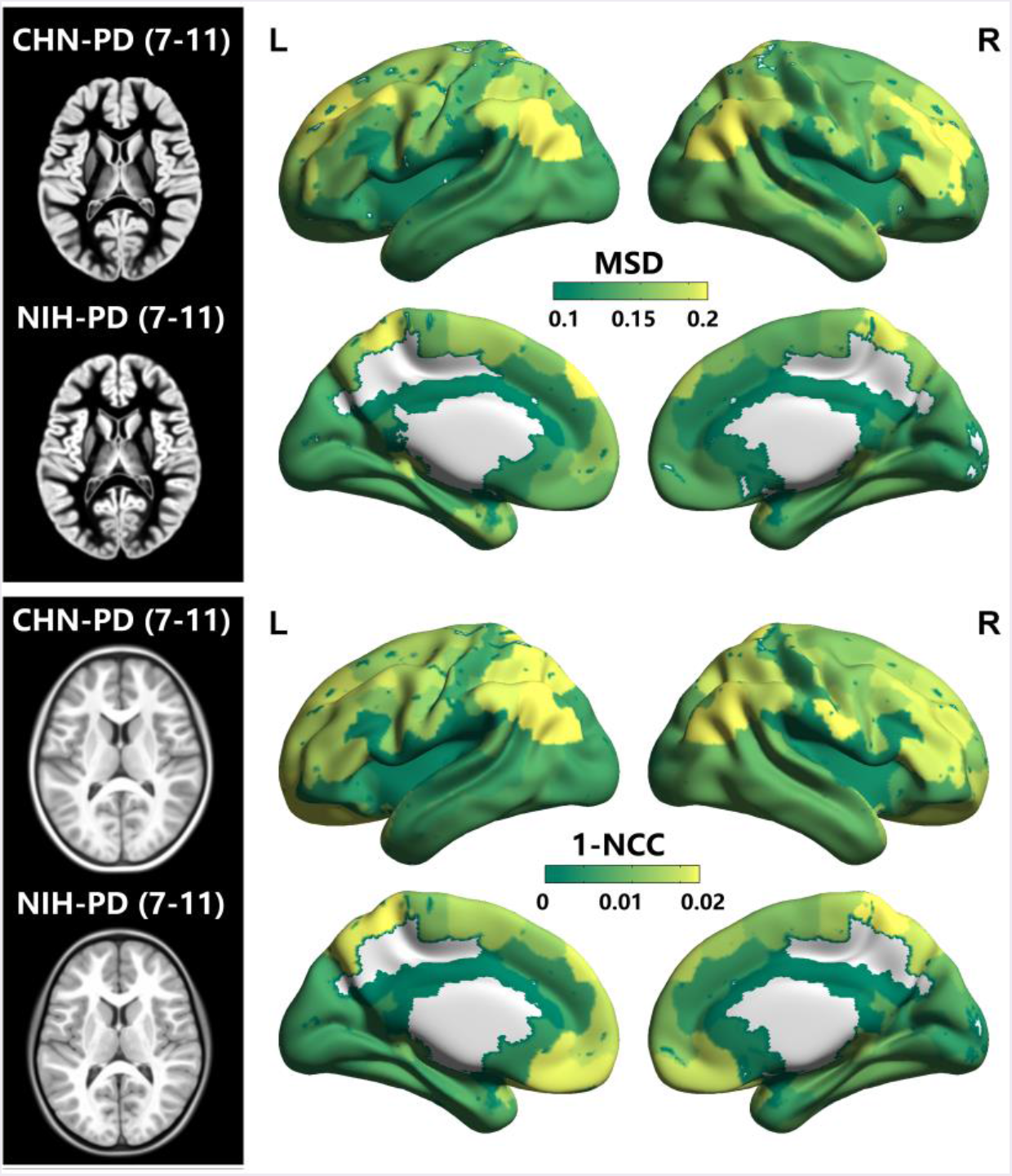
Regional anatomical differences between the CHN-PD templates and the NIH-PD templates. The detailed slices of the age and gender-matched Chinese pediatric brain template and the NIH pediatric brain template are given on the left (upper panel: gray matter probability map; lower panel: T1-weighted brain template). Distributions of regional anatomical differences from the perspective of the mean square difference (MSD) and the normalized cross correlation (NCC) are shown in a 3D surface view, with colors from green to yellow coding the index value from low to high. Regions with relatively high anatomical differences are mainly located in the bilateral angular gyrus and supramarginal gyrus, bilateral dorsolateral prefrontal and inferior frontal cortexes, and bilateral somatosensory cortex in both indexes. The visualization of the 3D surface view was accomplished using the BrainNet Viewer software (http://www.nitrc.org/projects/bnv/) (Xia, et al., 2013).

**Table 3.** Regions showing high anatomical differences between the CHN-PD and NIH-PD templates

| Regions | MSD | Regions | 1-NCC |
| --- | --- | --- | --- |
| Left angular gyrus, Wernicke's area | 0.224 | Right somatosensory cortex | 0.022 |
| Right dorsolateral prefrontal cortex | 0.222 | Right primary somatosensory cortex | 0.02 |
| Left dorsolateral prefrontal cortex | 0.222 | Left angular gyrus, Wernicke's area | 0.017 |
| Right angular gyrus, Wernicke's area | 0.221 | Left somatosensory cortex | 0.017 |
| Left somatosensory cortex | 0.218 | Right angular gyrus, Wernicke's area | 0.016 |
| Right primary, somatosensory cortex | 0.214 | Right primary gustatory cortex | 0.016 |
| Right somatosensory cortex | 0.212 | Left visuo-motor coordination | 0.015 |
| Left supramarginal, Wernicke's area | 0.204 | Left orbitofrontal, rostral superior frontal gyrus | 0.015 |
| Left dorsolateral prefrontal cortex | 0.204 | Right dorsolateral prefrontal cortex | 0.014 |
| Right parstriangularis, Broca's area | 0.203 | Left dorsolateral, prefrontal cortex | 0.014 |
| Right dorsolateral, prefrontal cortex | 0.201 | Left supramarginal, Wernicke's area | 0.014 |
| Right supramarginal, Wernicke's area | 0.2 | Right parstriangularis, Broca's area | 0.014 |
| Left piriform cortex | 0.199 | Right orbitofrontal, rostral superior frontal gyrus | 0.014 |
|  |  | Right visuo-motor coordination | 0.013 |
|  |  | Right supramarginal, Wernicke's area | 0.013 |

Regions with indexes greater than "mean+std" are shown and listed in descending order in the table.

### Different prediction power of brain age in Chinese pediatrics using the CHN-PD and the NIH-PD atlases

Using the HLG dataset (Dataset 2) as the training set and the healthy ADHD200 samples from Beijing site (Dataset 3) as the test set, a higher accuracy of age prediction was obtained by employing the CHN-PD brain template (with the highest correlation r = 0.40) during image normalization than by employing the NIH-PD brain template (with the highest correlation r = 0.38) (Fig. 6A). In addition, we further exchanged the training and test samples to perform a cross-validation of the prediction. A higher accuracy was also obtained via the implementation of the CHN-PD brain template (with the highest correlation r = 0.48) than with the NIH-PD brain template (with the highest correlation r = 0.43) (Fig. 6B). The better performance of the Chinese pediatric brain template was reproducible in different feature-preprocessing operations and different regression models (Table 4).

**Figure 6.**
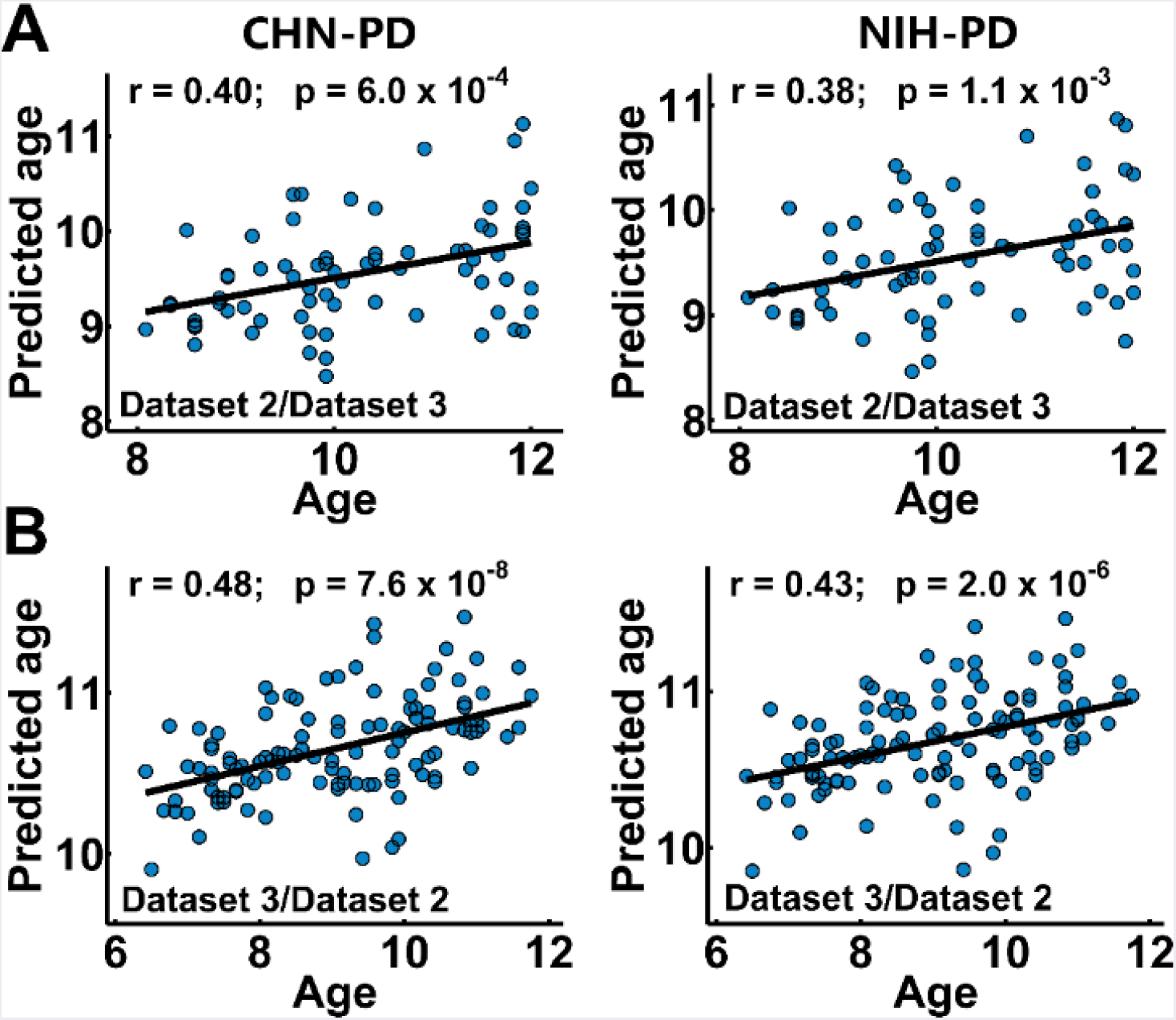
The template effect on the accuracy of age prediction. Pearson correlation coefficients between the actual and predicted ages were calculated to represent the prediction accuracy. By using the healthy Chinese children in Dataset 2 and Dataset 3 as the training/test samples (marked on the bottom of each figure) alternatively (A and B separately), the prediction framework employing the Chinese pediatric template (CHN-PD) as the normalization target consistently showed a higher prediction accuracy for brain age than the use of the NIH pediatric (NIH-PD) template.

**Table 4.**
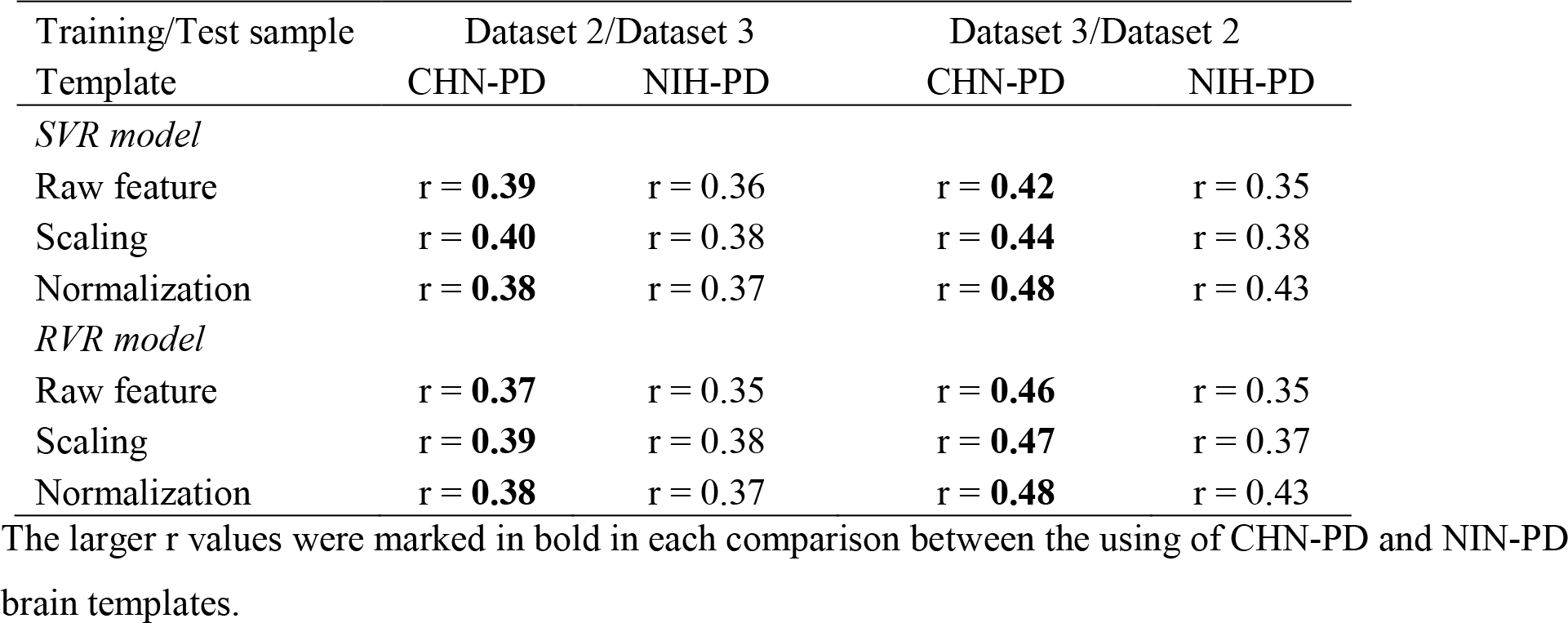
The accuracy of age prediction in different regression frameworks by employing two brain templates as the normalizing target

## Discussion

In the current study, we constructed a set of Chinese age-appropriate brain atlases using a large sample of high quality MRI images of Chinese children. These Chinese pediatric (CHN-PD) atlases included *head/brain*, symmetric and gender-specific MRI templates aged from 6-12 years old and with one-year intervals (Figs. 3 and 4). The proposed Chinese pediatric templates showed obvious anatomical differences in the lateral frontal and parietal cortex regions, somatosensory cortex and language-related areas compared with the NIH-PD templates. In the Chinese pediatric datasets, we found higher age prediction accuracy by using the Chinese pediatric brain template as the normalization target than by using the NIH pediatric brain template.

Two previous studies have made efforts to build Chinese pediatric brain atlases (Luo, et al., 2014; Xie, et al., 2015). However, there are several major differences between our work and these two earlier studies. First, our MRI atlases (Fig. 3 and 4) employed high-quality images with high signal noise ratio obtained by the advanced, state-of-the-art 3.0T Prisma scanners; in contrast, the two earlier studies used brain images obtained in 1.5T MRI scanner (Luo, et al., 2014) and 3.0T Trio scanner (Xie, et al., 2015). Second, our atlases included the relatively larger sample size (N = 328) than the two earlier studies (N = 53 and 138), which enabled refined one-year intervals among the pediatric MRI templates. This refinement is important for obtaining an accurate description for the elaborate growth trajectories of children brain during a period with rapid and dynamic structural and functional changes (Cao, et al., 2016; Giedd, et al., 1999; Lebel and Beaulieu, 2011; Levman, et al., 2017; Sowell, et al., 2003; Walhovd, et al., 2017). Third, unlike the rigid transformation used in the two previous studies (Luo, et al., 2014; Xie, et al., 2015), we applied a nine-parameters affine transformation on the individual images to the initial ICBM152 brain target, which reduced the differences in the circumference between Chinese and Western brain while introduced more detailed and compatible regional features of the Chinese pediatric templates. Fourth, the consistent coordinate system according to the MNI space of our atlases makes the application of our templates convenient and generalizable. Finally, our brain atlases included symmetric and gender-specific types, which were not included in the two earlier studies (Luo, et al., 2014; Xie, et al., 2015).

Notably, we used an unbiased model construction algorithm for the construction of brain atlases. It has several methodological advantages and has been widely used in the creation of MRI templates (De Leener, et al., 2018; Fonov, et al., 2011; Fonov, et al., 2009; Fonov, et al., 2014). The hierarchical local nonlinear registrations were iteratively corrected by the common features of the population in the construction process, which can obtain abundant age-related features while maintaining clear and sharp averaged tissue contrast at the same time. Rather than the usual linear interpolation in the image resampling, the cubic spline interpolation employed in the current pipeline can yield slightly better results (Thévenaz, et al., 2000). Another widely used template construction strategy is the diffeomorphic framework (Avants, et al., 2009), which can guarantee a smooth and differentiable nonlinear transformation during model iterations. We did not choose this framework for two reasons. First, a recent study has shown that the employment of a fully diffeomorphic algorithm may not automatically guarantee an increase in accuracy during template construction (Fonov and Collins, 2018). Second, the cortical folding pattern of the brain template could be altered when using different atlas construction methods. To make an accurate comparison between the CHN-PD templates and the widely used NIH-PD templates, a consistent template construction framework is used.

Several studies have demonstrated significant differences in overall brain morphology (e.g., size, shape and volume) between the Chinese and Western brain templates (Tang, et al., 2010; Xie, et al., 2015). Chinese children brain templates are generally shorter, wider, and taller than the age-appropriate American templates (Xie, et al., 2015). Our study further extended these structural differences to a detailed regional level (Fig. 5). The regions showed obvious anatomical differences that were mainly located in the sensorimotor regions and several high-order function regions such as the dorsal attention and language-related regions (e.g., Broca’s and Wernicke’s areas). These regional differences were quite similar to the previous multi-cultural brain studies in which Chinese adults showed significantly thinner cortical thickness in the premotor cortex, inferior frontal gyrus and supramarginal gyrus (Chee, et al., 2011); smaller cortical volume; and larger surface areas in the bilateral superior and medial prefrontal and the bilateral orbitofrontal gyrus (Tang, et al., 2018) compared with Western. These consistent regional differences are reasonable because the high-order functions related to cultural and educational factors such as the language abilities have developed prominently in children at school age. Our results (Fig. 5) provided a detailed brain map of the potential regional influences when adopting a dis-matched children brain template in Chinese pediatric MRI studies. However, how these differences affect the assessments on brain development in Chinese pediatric cohorts or act on the estimations of multi-cultural effects on child brain still needs further investigation.

An accurate prediction of the individual brain age is valuable for both typical and atypical children development investigations. Individual deviations towards the norm age trajectory of brain development may serve as potential markers for brain health and disorders (Cole and Franke, 2017; Dosenbach, et al., 2010). Studies have shown that brain structural images can be used to predict the individual age with high accuracies for both development (5-18 years old) and aging (19-86 years old) populations (Franke, et al., 2012; Franke, et al., 2010). A recent MRI study has investigated several methodological factors during image feature generation in order to improve the accuracy of age prediction (Monté-Rubio, et al., 2018). Our results further revealed that the application of Chinese specific brain templates can help to facilitate the accurate prediction of brain age in Chinese pediatrics (Fig. 6). Our prediction accuracy did not reach a high value compared with other studies. However, its performance was acceptable since the age range of the participants in the current study is relatively narrow (Dataset 2: 6–12 years old; Dataset 3: 8–12 years old) and the sample used for training the model is relatively small (Dataset 2: 114 subjects; Dataset 3: 71 subjects), which may increase the difficulty of prediction (Cui and Gong, 2018). Although the aim of the current study was not to seek the highest prediction accuracy for age, our results indeed indicated that under the same analysis framework, the Chinese pediatric brain template could increase the accurate estimation of age effects in the Chinese children population.

Several issues need be further addressed in our study. First, although the sample size used for the template construction has been improved, more subjects would be beneficial for sure. Second, future studies could make efforts towards developing novel, cohort-specific brain templates using different sub-populations (such as younger children or disease-related atlases) to obtain a comprehensive representation of Chinese pediatrics. Third, multi-atlas libraries containing multi-modality templates such as white matter atlases based on diffusion weighted images should be established to provide abundant brain contrasts. Finally, we anticipated that the proposed Chinese children brain atlases can be used for future studies on typical and atypical development in Chinese pediatric populations, which may bring a better understanding of the development of pediatric population in China.

## Acknowledgements

This work was supported by the Changjiang Scholar Professorship Award (T2015027), and the National Natural Science Foundation of China (81620108016, 31521063, 31522028), the Fundamental Research Funds for the Central Universities (2017XTCX04),

